# ComHub: Community predictions of hubs in gene regulatory networks

**DOI:** 10.1101/840959

**Authors:** Julia Åkesson, Zelmina Lubovac-Pilav, Rasmus Magnusson, Mika Gustafsson

## Abstract

**Summary:** Hub transcription factors, regulating many target genes in gene regulatory networks (GRNs), play important roles as disease regulators and potential drug targets. However, while numerous methods have been developed to predict individual regulator-gene interactions from gene expression data, few methods focus on inferring these hubs. We have developed ComHub, a tool to predict hubs in GRNs. ComHub makes a community prediction of hubs by averaging over predictions by a compendium of network inference methods. Benchmarking ComHub to the DREAM5 challenge data and an independent data set of human gene expression, proved a robust performance of ComHub over all data sets. Lastly, we implemented ComHub to work with both predefined networks and to do standard network inference, which we believe will make it generally applicable.

**Availability:** Code is available at https://gitlab.com/Gustafsson-lab/comhub

## 1 Introduction

As of today, there is a plethora of methods available to reverse engineer gene regulatory networks (GRNs) from gene expression data [Haury *et al*., 2012, Faith *et al*., 2007, Margolin *et al*., 2006, Huynh-Thu *et al*., 2010]. Correspondingly, there have been numerous benchmark studies to evaluate the ability of these methods to correctly predict known networks, culminating in the DREAM5 challenge [Marbach *et al*., 2012]. In the DREAM5 challenge, the main conclusion was that combining multiple GRN prediction tools gives the most robust and accurate performance. GRN inference has often been employed to find genes with a disproportionately large amount of connections to the rest of the network, henceforth referred to as hubs, [Gustafsson *et al*., 2015, Alvarez *et al*., 2018], with some tools being designed especially for extracting these hubs from mRNA expression data [Lefebvre *et al*., 2010, Liu *et al*., 2011]. However, few studies, if any, have analysed the benefit of combining inference methods when inferring hubs, such as done with GRNs in [Marbach *et al*., 2012]. The field is also lacking software that makes a community prediction of hubs. To this end we built ComHub, a method and accompanying software that combines several GRN inference methods, and thereafter ranks hubs in order of centrality measured by the outdegree of each gene. We first applied ComHub to the DREAM5 challenge, and observed hub rankings that were robustly outperforming almost all the ingoing DREAM5 GRN inference methods in terms of correlation between predicted and gold standard outdegrees. Furthermore, we validated this performance using human gene expression data. In summary, ComHub provides an open source Python software to infer hubs from expression data by compiling GRNs in a novel way.

## 2 Software description

We implemented ComHub as a Python 3 package that takes predicted GRNs as input, and from these GRNs ranks each regulator by average outdegree. If no a priori putative networks exist, ComHub takes two text files as input, one containing gene expression data and one with a list of potential gene expression regulators, e.g. transcription factors. ComHub utilizes widely used GRN inference methods, and to capture properties across all types of approaches, we sough to include default methods of a broad set of inference classes. From the field of linear regression, we implemented a bootstrap ElasticNet [Zou *et al*., 2005], and the “trustful inference of gene regulation with stability selection” (TIGRESS) [Haury *et al*., 2012]. Mutual information methods added were the “context likelihood of relatedness” (CLR) [Faith *et al*., 2007] and the “algorithm for the reconstruction of accurate cellular networks” (ARACNE) [Margolin *et al*., 2006]. We also incorporated the correlation-based method of calculating the absolute value of the Pearson correlation coefficient (PCC) [Butte *et al*., 2000], and the tree-based GRN inference method “gene network inference with ensemble of trees” (GENIE3) [Huynh-Thu *et al*., 2010]. To not limit ComHub to the default methods, we also added the option for the user to directly input GRNs of any chosen inference method.

ComHub operates in mainly three steps. First, if not provided by the user, a set of putative GRNs, containing confidence-ranked gene-gene interactions is created from the ingoing methods. Second, the correlations of hub rankings between these predefined GRNs are calculated, and a threshold of edges that gives the most similar consensus is imposed on the ranked gene-gene interactions. Third, the average method outdegree of each regulator is calculated (Fig. 1).

**Figure 1:**
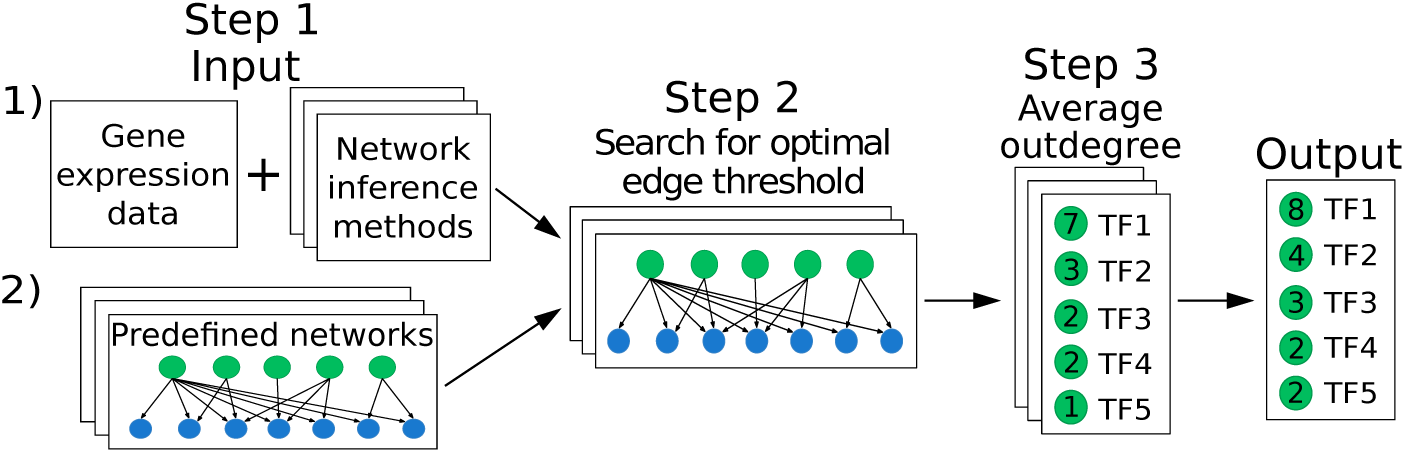
The workflow of ComHub. ComHub either takes as input 1) Gene expression data and applies a set of network inference methods or 2) predefined networks. ComHub identifies an optimal edge threshold for the predicted GRNs and the outdegrees of each regulator is averaged over the method predictions.

## 3 Results

### 3.1 An intra-community assessment identifies better edge prediction thresholds

We developed ComHub by implementing the approach on predictions of the 35 network inference methods in the DREAM5 challenge, where networks were reverse engineered from *Escherichia coli*, and *in silico* gene expression data. Each of the 35 GRNs contained 100,000 ranked edges, and to choose a cut-off we compared the similarity between GRNs as a function of included edges. We observed that the peak of this similarity corresponded to a peak in performance (Fig. S1). This finding is possibly applicable for all methods aiming to construct consensus GRNs, as it gives an independent approach to estimate information in data.

### 3.2 ComHub robustly predicts hub TFs

As noted by [Marbach *et al*., 2012], GRN inference methods have an overall diverse performance depending on data properties. This variation also applies to the ability to predict hubs, as measured by gene outdegree (Fig. S2b). To address this limitation, we applied ComHub to construct a community hub prediction. By applying ComHub to sets of randomly drawn networks from the DREAM5 contestants we observed a robust network, converging with the number of in-going methods (Fig. S2a). Notably, this convergence occurs at a relatively low number of in-going DREAM5 predictions, and at only 6 methods the average PCC measures 85% and 90% of the maximal PCC for *E. coli* and *in silico*, respectively. When including all 35 methods, we observed a PCC between the predicted and gold standard outdegree of 0.38 and 0.71 for *E. coli* and *in silico* (Fig. S2). In addition, ComHub outperformed the community approach presented in the DREAM5 challenge, where the consensus was taken on each gene-gene interaction individually, on *E. coli* (Fig. S2).

### 3.3 ComHub was validated on human RNA-Seq data

To verify the applicability of ComHub on an independent dataset, we used a compendium of human gene expression data across tissues from the Human Protein Atlas (HPA) ([Uhlén *et al*., 2015]), along with the human protein-protein interaction network from STRINGdb as a gold standard. First, ComHub identified the number of interactions to be used in the predicted network to 100,000 by assessing the similarity between the methods (Fig. S3a). Again, this point coincided with the peak in performance for individual methods. We observed ComHub to again outperform the community of individual edges, as presented in the DREAM5 challenge analysis (Fig. S3b,c). Lastly, we assessed the overall performance of ComHub and the six individual methods on the HPA, *E. coli*, and *in silico* data sets, and noted that ComHub outperforms the other methods (Fig S4).

## 4 Conclusions

ComHub is a novel tool that assembles knowledge derived from common hub inference methods to improve hub predictions. This tool will benefit the research on hub-based biomarker discovery, which can contribute to understanding of complex disease.

## References

Alvarez, M.J. et al. (2018) A precision oncology approach to the pharmacological targeting of mechanistic dependencies in neuroendocrine tumors. Nat. Genet., 50, 979–989.

Butte, A.J. and Kohane, I.S. (2000) Mutual Information Relevance Networks: Functional Genomic Clustering Using Pairwise Entropy Measurements. Pacific Symp. Biocomput., 5, 415–426.

Faith, J.J. et al. (2007) Large-scale mapping and validation of Escherichia coli transcriptional regulation from a compendium of expression profiles. PLoS Biol., 5, 0054–0066.

Gustafsson, M. et al. (2015) A validated gene regulatory network and GWAS identifies early regulators of T cell-associated diseases. Sci. Transl. Med., 7, 1–10.

Haury, A.C. et al. (2012) TIGRESS: Trustful Inference of Gene REgulation using Stability Selection. BMC Syst. Biol., 6, 1–17.

Huynh-Thu, V.A. et al. (2010) Inferring regulatory networks from expression data using tree-based methods. PLoS One, 5, 1–10.

Lefebvre, C. et al. (2010) A human B-cell interactome identifies MYB and FOXM1 as master regulators of proliferation in germinal centers. Mol. Syst. Biol., 6, 1–10.

Liu, Q. and Ihler, A. (2011) Learning scale free networks by reweighted l1 regularization. J. Mach. Learn. Res., 15, 40–48.

Marbach, D. et al. (2012) Wisdom of crowds for robust gene network inference. Nat. Methods, 9, 796–804.

Margolin, A.A. et al. (2006) ARACNE: An algorithm for the reconstruction of gene regulatory networks in a mammalian cellular context. BMC Bioinformatics, 7 (Suppl1), 1–15.

Uhlén, M. et al. (2015) Tissue-based map of the human proteome. Science, 347, 1260419–1260419.

Zou, H. and Hastie, T. (2005) Regularization and variable selection via the elastic-net. J. R. Stat. Soc., 67, 301–320.

