## Supplementary material for "ComHub: Community predictions of hubs in gene regulatory networks"

### 1 ComHub

#### 1.1 ComHub Workflow

ComHub is a tool for predicting network hubs in gene regulatory networks (GRNs). ComHub makes hub predictions by averaging regulator outdegrees over a compendium of reverse engineered GRNs. ComHub operates in three steps:

**Step 1:** ComHub has a set of network inference methods implemented to infer GRNs from gene expression data (see section 1.2). As an option ComHub takes GRN predictions directly as input. It is up to the user to decide how many default methods and how many of the users own GRN predictions that should be used. The GRN predictions should be a tab separated text file, containing regulator-target interactions as the first two columns, any additional columns are not used. The regulator-target interactions should be ranked on a method specific confidence score.

**Step 2:** A threshold is set on the number of edges to include from the GRN predictions. ComHub searches for an optimal edge threshold by assessing the Pearson correlation coefficient (PCC) between each pair of method predictions over a range of edge thresholds (by default 1000 to 100,000 edges). The correlation is assessed between the regulator outdegrees for each edge threshold. The optimal edge threshold is selected where the correlation between method predictions (M) is maximised according to:

$$edgethreshold = \max_{1000 \leq e \leq 100,000} \frac{\sum_{i,j} PCC(M_i(e), M_j(e))}{C},$$

where (i,j) is all possible combinations of method predictions {(1,2),(1,3),...,(N-1,N)}, N is the number of method predictions, C is the number of possible combinations of method predictions (C = (N-1)N/2), and e is the edge threshold. The top-ranked regulator-target interactions passing the edge threshold are included in the networks.

**Step 3:** For the optimal edge threshold the outdegree of each regulator is averaged over all method predictions. The outdegree of each regulator is determined by counting all edges passing the set threshold. This is done for each predicted network separately, before calculating a combined score for each regulator according to:

$$score_k = \frac{1}{N}\sum_{l=1}^{N} outdegree_{kl}, k \in \{1, R\}, l \in \{1, N\},$$

where N is the number of method predictions and R is the number of regulators. The output of ComHub is a list of regulators ranked on the community outdegree ($score_k$).

#### 1.2 Network inference methods implemented

If no predefined networks are provided, ComHub relies on the assumption that applying methods using different types of approaches will result in more reliable predictions. ComHub has 6 default methods implemented: the linear regression methods TIGRESS and Elastic Net with bootstrap, the mutual information (MI) methods CLR and ARACNE, the correlation method absolute value of the PCC, and the tree-based method GENIE3 (table S1). All methods operates with 2 text files as input: one containing gene expression data and one with possible regulators. The output is a text file containing the inferred GRN with regulator-target interactions and accompanying confidence score. The regulator-target interactions are ranked on the confidence score. It is possible for the user to decide how many top ranked regulator-target interactions that should maximally be inferred by each method (by default 100,000 regulator-target interactions are maximally inferred).

| **Method** | **Description** | **Reference** |
|---|---|---|
| Bootstrap Elastic Net | 1) Elastic Net 2) bootstrapping | [1] |
| TIGRESS "trustful inference of gene regulation with stability selection" | 1) Least angle regression (LARS), 2) stability selection | [2] |
| CLR "context likelihood of relatedness" | 1) Computes a mutual information score for each regulator-target interaction 2) Filters interactions not significantly above the "background" distribution of MI scores. | [3] |
| ARACNE "algorithm for the reconstruction of accurate cellular networks" | 1) Computes a mutual information score for each regulator-target interaction 2) Applies data processing inequality to remove indirect interactions. | [4] |
| Absolute value of the PCC | Regulator-target interactions are ranked on the absolute value of the PCC between regulator-target gene expression. | [5] |
| GENIE3 "gene network inference with ensemble of trees" | Decomposes the network inference into different feature selection problems and applies tree-based ensemble methods on each sub-problem. | [6] |

Table 1. The network inference methods implemented in ComHub.

### 2 Benchmark

#### 2.1 DREAM5 data sets

ComHub were benchmarked on the *E. coli* and *in silico* data sets presented in the DREAM5 challenge [7]. In the DREAM5 challenge, contestants were given gene expression data from biological and *in silico* generated networks, with instructions to predict gene-to-gene interactions. The DREAM5 challenge included 35 network inference methods and one community network approach combining the results of the 35 network inference methods. The network inference methods were divided into 5 groups: Regression, MI, Correlation, Bayesian networks, Other approaches and Meta predictors. Each of the 35 contestants delivered a maximum of 100,000 ranked regulator-target interactions. For each of the *E. coli* and *in silico* data set gold standards were used to evaluate the performance. A more detailed description of network inference methods, gene expression data, and gold standards can be found in [7].

#### 2.2 Performance assessment of the DREAM5 methods

We evaluated the 35 network inference methods and the community network approach presented in the DREAM5 challenge based on how well the methods predicted hubs in the *E. coli*, and *in silico* networks. The performance were assessed by calculating the PCC between regulator outdegrees of each method prediction and gold standards. We used the gold standards compiled during the DREAM5 challenge. The regulator outdegree is largely influenced by the number of inferred edges. Therefore, we first sought an independent measure to identify a threshold for edge inclusion.

#### 2.3 Identifying an edge threshold

To identify an edge threshold we assessed the correlation of regulator outdegree between each pair of predicted networks as a function of included interactions. Notably, the pairwise correlations corresponded well with the performance of the methods (Fig. S1), with the peak pairwise correlation coinciding with the optimal performance. This attribute allows for an unbiased approach to estimate the number of edges to be included in the GRN prediction. Using this criterion, we estimated the optimal number of included edges to be 80,000 and 2000 for the *E. coli* and *in silico* data-sets, respectively. Using the gold standard network, we found the true optimal edge number to be 80,000 and 1000, with optimal performance either being correctly identified (*E. coli*), or within 2% of the optimal performance for *in silico*. Hence, the pairwise correlation between methods can be used as a measurement on how many inferred interactions that should be included in a community prediction. As noted by Marbach et al. [7], GRN inference methods have an overall diverse performance depending on data set, with different methods being the top-performer between the *E. coli* and *in silico* data sets. We found this variation to also be reflected in the ability to predict hubs (Fig S2b). To address this limitation, we applied ComHub to construct a community hub prediction.

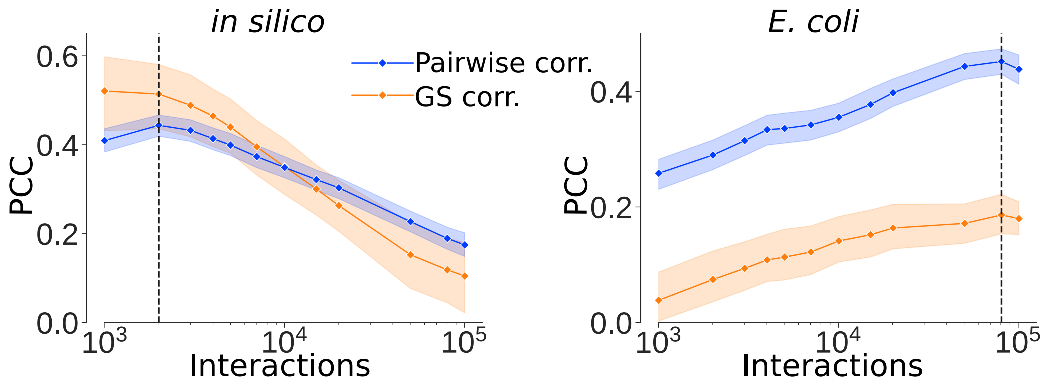

Figure 1: The average pairwise correlation, assessed with Pearson correlation coefficient (PCC), between method predictions (blue) compared to the average method performance (orange) for *in silico* and *E. coli* data sets. The dotted lines shows were the optimal edge threshold were selected.

#### 2.4 In-going methods effect on the performance of ComHub

ComHub were applied to sets of randomly drawn predicted networks from the DREAM5 contestants. The networks were randomly drawn using two different approaches. The networks were randomly drawn with replacement from 1) the total set of 35 predicted networks (random hub community), and 2) the set of 35 predicted networks weighted on group of method, resulting in the combinations being balanced between the groups (ComHub). We assessed the average performance of ComHub as a function of the number of in-going predictions (Fig. S2a). We observed robust hub predictions converging with the number of in-going methods. At only 6 methods the average PCC measured 85% and 90% of the maximal PCC (combining 35 method predictions) for *E. coli* and *in silico*, respectively. Hence, a relatively low number of network inference methods were enough to make robust predictions. As a result we implemented 6 default methods in ComHub. In addition, combining methods with different approaches seemed to have a positive effect on the performance. Therefore, the ComHub default methods covers several different approaches. We compared the performance of ComHub with the DREAM5 community network approach, and observed similar behaviour. Notably, ComHub performed considerably better than the DREAM5 community network approach, in the case of the biologically derived network.

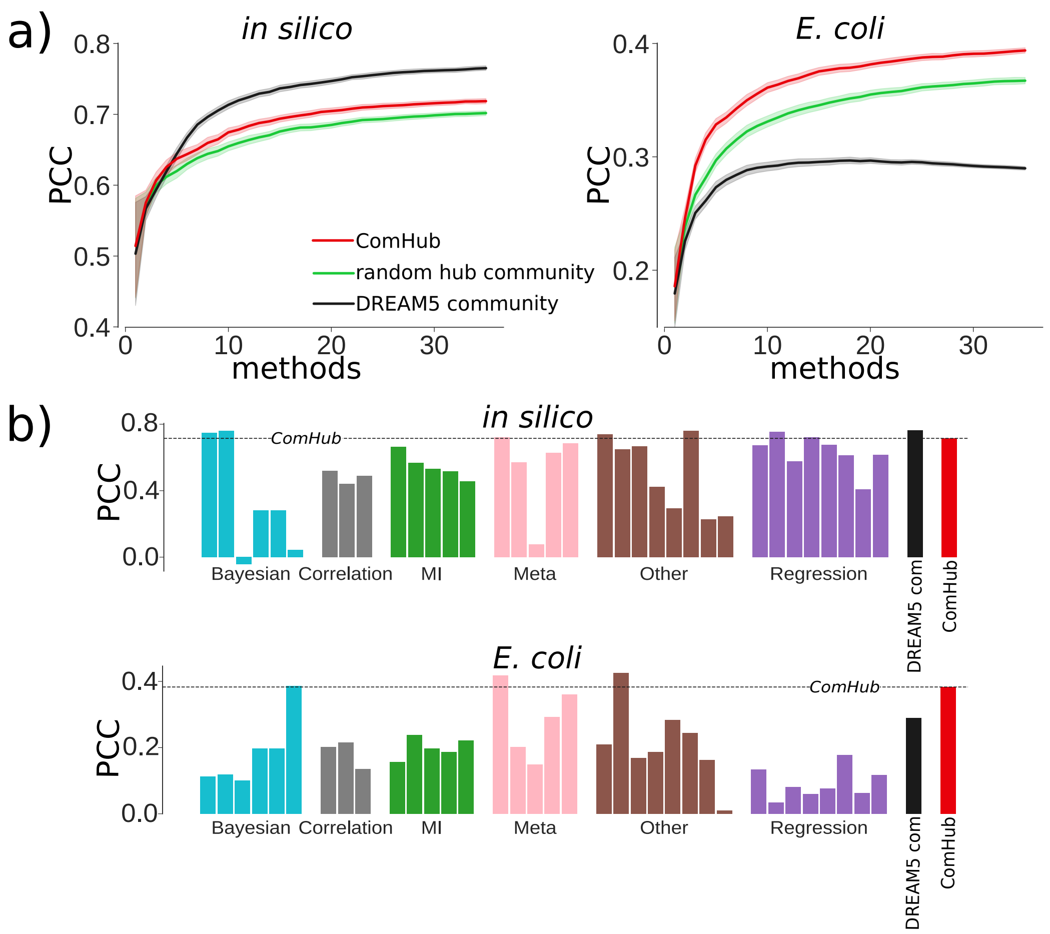

Figure 2: a) The performance of ComHub (red) with combinations weighted on method approach, the random hub community (green) and the DREAM5 community network (black) as a function of in-going methods. b) The maximal performance (combining 35 method predictions) of ComHub, DREAM5 network approach and each of the DREAM5 participant method, on the *in silico* and *E. coli* data sets.

#### 2.5 ComHub performance on a human gene expression data set

ComHub were applied on gene expression data from 37 tissues and 64 cell lines obtained from the Human Protein Atlas (HPA) [8]. A list of possible transcription factors (TFs) were obtained from DBD: Transcription factor prediction database [9]. As a gold standard we used a protein-protein interaction network obtained from STRINGdb version 10.5 [10], where edges with a confidence score>700 were included. The gold standard only contained TFs and target genes present in the gene expression data and in the list with possible TFs.

First, ComHub identified the number of interactions to be used in the predicted network to 100,000 by assessing the pairwise correlation between the methods (Fig. S3a). Strikingly, the point of 100,000 included edges again coincided with the optima for individual methods' ability to predict hubs. When using the 100,000 top ranked edges to construct a community prediction, we again observed the effect of including more GRN inference algorithms to saturate at 3 methods (Fig. S3b). Moreover, we observed ComHub to again outperform the community of individual edges, as presented in the DREAM5 challenge analysis (Fig. S3b-c).

Next, we studied the biological interpretation of the ComHub prediction on the HPA data set. We found the community hub prediction to be non-linear, with the top 1% TFs having >10% of the identified interactions. Among those master regulators we found TFs such as FOXM1, YBX1 and SMARCC1 which are related to different types of carcinoma.

Lastly, we assessed the overall performance of ComHub, the six individual methods, and the DREAM5 community approach on the HPA, *E. coli*, and *in silico* data sets. A combined score were computed for each method ($\mathrm{M_i}$) according to:

$$combinedscore(M_i) = \frac{PCC_{insilico}(M_i)}{max(PCC_{insilico})} + \frac{PCC_{E.coli}(M_i)}{max(PCC_{E.coli})} + \frac{PCC_{HPA}(M_i)}{max(PCC_{HPA})}$$

Notably, ComHub outperforms the six individual methods and the DREAM5 community approach (Fig. S4).

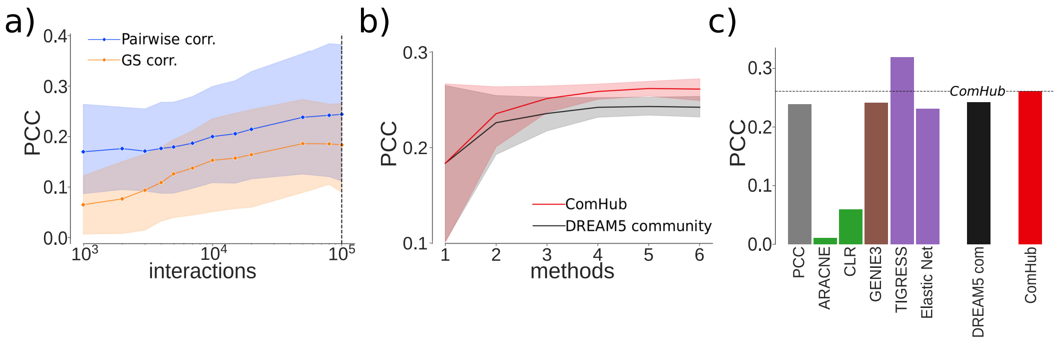

Figure 3: The performance of ComHub on the human gene expression data set. a) The pairwise correlation between the method predictions (blue), compared to average method performance (orange). The dotted line shows where the edge threshold were selected. b) The performance of ComHub (red) and the DREAM5 community network approach (black) as a function of in-going methods. c) The performance of ComHub, the DREAM5 community network approach and each of the 6 network inference methods used.

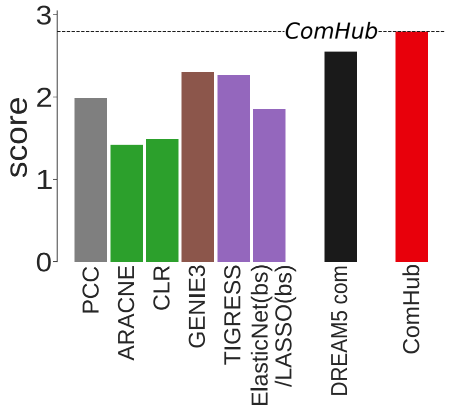

Figure 4: The combined performance of ComHub, the DREAM5 community approach and the six individual methods on the HPA, *E. coli*, and *in silico* data sets

### References

[1] Zou,H. and Hastie,T. (2005) Regularization and variable selection via the elastic-net. *J. R. Stat. Soc.*, **67**, 301–320.

[2] Haury,A.C. et al. (2012) TIGRESS: Trustful Inference of Gene REgulation using Stability Selection. *BMC Syst. Biol.*, **6**, 1–17.

[3] Faith,J.J. et al. (2007) Large-scale mapping and validation of Escherichia coli transcriptional regulation from a compendium of expression profiles. *PLoS Biol.*, **5**, 0054–0066.

[4] Margolin,A.A. et al. (2006) ARACNE: An algorithm for the reconstruction of gene regulatory networks in a mammalian cellular context. *BMC Bioinformatics*, **7**(Suppl1), 1–15.

[5] Butte,A.J. and Kohane,I.S. (2000) Mutual Information Relevance Networks: Functional Genomic Clustering Using Pairwise Entropy Measurements. *Pacific Symp. Biocomput.*, **5**, 415–426.

[6] Huynh-Thu,V.A. et al. (2010) Inferring regulatory networks from expression data using tree-based methods. *PLoS One*, **5**, 1–10.

[7] Marbach,D. et al. (2012) Wisdom of crowds for robust gene network inference. *Nat. Methods*, **9**, 796–804.

[8] Uhlén,M. et al. (2015) Tissue-based map of the human proteome. *Science*, **347**, 1260419–1260419.

[9] Wilson,D. et al. (2008) DBD - Taxonomically broad transcription factor predictions: New content and functionality. *Nucleic Acids Res.*, **36**, 88–92.

[10] Szklarczyk,D. et al. (2015) STRING v10: Protein-protein interaction networks, integrated over the tree of life. *Nucleic Acids Res.*, **43**, D447–D452.
